# Multigenerational machine learning-based genomic prediction for dermo resistance in eastern oyster *Crassostrea virginica*

**DOI:** 10.64898/2026.09.08.750100

**Authors:** Henry Sun, Paul Coyne, Zhenwei Wang, Sandra Casas, Jerome La Peyre, Mason L. Williams, Scott Rikard, David Bushek, Juliet Wong, Ximing Guo

## Abstract

Dermo disease caused by the protist *Perkinsus marinus* poses a major threat to Eastern oyster aquaculture. We previously conducted genomic selection for dermo resistance and found increased effectiveness over phenotypic selection. Here, we report improved genomic predictions using combined data from three successive generations. We evaluated nine different genomic selection models, including three supervised machine learning architectures: gradient boosting, logistic regression, and random forest. Combining data across multiple generations did not, by itself, substantially improve the accuracy of most models, but the increased sample size provided genotyping confidence of loci with rare alleles. Correlation accuracy of all models significantly increased with the inclusion of low-frequency variants and strong-effect markers identified through a genome-wide association study. Gradient boosting machine learning models outperformed other genomic selection models across all training scenarios, suggesting enhanced capacity to learn generalizable genomic signals associated with dermo resistance. The best gradient boosting model achieved a peak correlation accuracy of 0.410, a substantial improvement over the previous peak accuracy of 0.274 from traditional models. Together, our results highlight the potential of machine learning for genomic selection and the significance of rare variants in determining dermo resistance.

## Introduction

The Eastern oyster (*Crassostrea virginica*) is a species of critical ecological and economic importance, with the aquaculture production in the United States in 2024 valued at over $195 million [01]. As filter-feeders and habitat engineers, Eastern oysters provide essential ecosystem services and contribute to the protection of the coastline against erosion [02, 03, 04]. Despite these benefits, Eastern oyster populations have experienced large declines driven by multiple stressors associated with overharvesting, habitat degradation, disease, and climate change [05, 06]. Diseases are persistent stressors limiting Eastern oyster reef resilience, recovery, and aquaculture development [07]. One of the most consequential diseases affecting Eastern oysters is dermo, caused by the protist parasite *Perkinsus marinus*. *Perkinsus marinus* infects filter-feeding oysters through the water column, proliferating within host tissues and impairing physiological function [08]. Dermo has historically inflicted yearly mortalities that can exceed 50% of marked-sized (>75 mm) oysters [9, 10, 11], and still poses a major threat to both natural reefs and aquaculture operations. As such, selection for dermo resistance has been a major priority of Eastern oyster breeding [12, 13, 14].

Selective breeding efforts against dermo in oysters have traditionally relied on phenotypic selection, in which individuals are chosen for spawning based on observable traits such as survival under disease pressure [15, 16]. However, dermo resistance has low to moderate heritability and is likely a polygenic trait governed by many small effect genes, with evidence of balancing selection between alleles beneficial for dermo resistance linked to alleles detrimental for larval survival [17, 18, 19]. The low heritability and polygenic nature of dermo resistance reduce the overall effectiveness of traditional phenotypic selective breeding, which is also slow because of the combination of the long oyster breeding cycle and dermo pathogenesis requiring 2-3 years.

Genomic selection (GS) is now widely used for selective breeding and genetic improvement of major agricultural crops and offers a promising alternative to phenotypic selection by leveraging genome-wide marker information to predict breeding values [20, 21]. Genomic selection captures the combined effects of many loci across the genome and can increase selective accuracy for polygenic traits governed by many small effect genes [22, 23]. Additionally, GS removes the need to wait multiple years to sufficiently phenotype oysters as disease-resistant, potentially greatly improving selective efficiency by genotyping and spawning oysters annually as soon as they reach reproductive maturity. The development of a high-density 66k single-nucleotide polymorphism (SNP) array for Eastern oysters has enabled the practical implementation of GS in breeding programs [24]. Genomic prediction models have been assessed for traits such as growth and low salinity tolerance in the Eastern oyster [25, 26]. Previously, we conducted GS for dermo resistance and demonstrated improved effectiveness over phenotypic selection [19]. Since then, we have conducted GS for two additional generations and accumulated a large multigenerational dataset for further analyses.

Despite the significance of dermo disease, we know little of its genetic mechanisms and genes determining resistance [17, 27]. The identification of dermo-resistant markers or genes can potentially enhance GS or provide candidate genes for editing. Genome-wide association studies (GWAS) with our previous single-generation dataset only identified loci of modest effects [19]. We recently performed GWAS on the large multigenerational dataset described in this study and identified novel candidate loci strongly associated with dermo resistance [28]. Previous studies have shown that GWAS can identify informative subsets of markers that improve genomic prediction accuracy when incorporated into GS models [29, 30, 31] highlighting the importance of evaluating the impact of GWAS-selected markers (GSMs) on genomic prediction accuracy.

Advances in machine learning (ML) have also introduced new approaches for GS [32, 33]. ML models have the potential to model complex, nonlinear relationships among genetic markers and improve prediction performance for complex traits [34]. Applications of ML models for GS in aquaculture species have shown promise [35, 36, 37], but ML model performance often varies greatly depending on the size and structure of training datasets [38]. Given these considerations, there is a need to evaluate ML approaches for genomic prediction in Eastern oysters, particularly in the context of important traits such as dermo resistance. However, direct comparisons between ML and conventional GS models for genomic predictions in Eastern oysters are missing.

In this study, we evaluated ML models for predicting dermo resistance using a large dataset from three successive generations of GS. By integrating data across generations, we increased the statistical power for GS and detection of rare variants. Using this combined dataset, we compared conventional and ML– based GS models and assessed the impact of incorporating rare variants and GSMs. Gradient boosting models showed the highest prediction accuracy, particularly following the inclusion of biologically relevant GSMs.

## Results

### Data quality control (QC)

We combined and analyzed genomic and phenotypic data from three generations of GS, with each training population comprising dead (susceptible) and live (resistant) oysters from lab-based dermo challenge experiments using the same protocol. The combined dataset contained 2,423 oysters genotyped at 65,893 SNPs with the 66K OysterCv SNP array [24]. After removing samples with < 90% individual call rate and SNP markers with a < 95% call rate and minor allele frequency (MAF) < 0.05, a total of 2,357 individuals and 39,923 SNPs remained, with 1,256 individuals (53.29%) surviving lab-based dermo challenge. There was a total of 493 samples from the F0 generation, 900 samples from the F1 generation, and 964 samples from the F2 generation, with 243 (49.29%), 519 (57.67%), and 494 (51.24%) survivors each, respectively.

### Genomic selection model performance across three generations

We evaluated the genomic prediction accuracy (hereafter referred to as correlation accuracy) of nine GS models. Correlation accuracy was defined as the Pearson correlation between model predictions of genome-estimated breeding values (GEBVs) with true binary phenotypes of survival against dermo challenge. The nine models included six conventional GS models, Bayesian B (BayesB), Bayesian Ridge Regression (BRR), genomic best linear unbiased prediction (GBLUP), extended GBLUP (EGBLUP), Bayesian least adjusted selection shrinkage operator (LASSO), and reproducing kernel hilbert space (RKHS), and three ML models, gradient boosting (GB), logistic regression (LR) and random forest (RF). BayesB, GB, LR, and RF are new models not previously tested in our published study of genomic selection models trained on the F0 generation [19].

To assess whether training on larger datasets improves model correlation accuracy, we first compared changes in accuracy between models trained on the entire dataset (n = 2357) and the most recent F2 generation (n = 964). At a MAF of 0.05, all conventional GS models except BayesB saw significant decreases in correlation accuracy ranging from 0.1% (GBLUP) to 6.4% (EGBLUP) when trained on the entire three generations of data (Supplementary Figure 1, Supplementary Table 2). Of the ML models, the RF model sharply decreased in performance by 18% (p = 1.08 * 10^-5^, Wilcoxon signed-rank test) on the entire dataset, while the LR model did not show a significant change in correlation accuracy (p = 0.971, Wilcoxon signed-rank test). Contrastingly, the GB and BayesB models exhibited a 3.3% (p = 0.0185, Wilcoxon signed-rank test) and 3.4% increase (p = 0.0185, Wilcoxon signed-rank test), respectively, when trained on the entire dataset compared to just the F2 generation.

### Model performance at lower MAF

We then assessed the influence of rare variants (loci with MAF < 0.05) on correlation accuracy by lowering the MAF QC threshold to 0.01 and 0.005. Following this adjusted QC, 48,143 and 50,279 SNPs were retained at MAF 0.01 and MAF 0.005 in the training dataset, respectively, with six additional individuals (total n = 2,363) passing the individual call rate QC at both MAF 0.01 and 0.005, two from the F0 generation and four from the F1 generation.

There were significant differences in model performances at all three MAF filter levels trained on both the F2 generation only and all three generations of data (all generations: MAF 0.05 Kruskal-Wallis p = 1.91 * 10^-13^, MAF 0.01 Kruskal-Wallis p = 6.9 * 10^-15^, MAF 0.005 Kruskal-Wallis p = 1.41 * 10^-14^) (Figure 1). The GB models had the highest overall correlation accuracy across all training dataset permutations, regardless of training dataset generation or MAF filter; across the six combinations of training datasets, the highest overall accuracy was achieved by the GB model trained on all three generations of data at MAF 0.01 (0.319). Here, we briefly summarize results from evaluation across all three generations of data combined. At MAF 0.05, the GB model had a 14% increased accuracy than the second-best LR model (Dunn test p = 0.19, Cohen’s d = 5.635) and a 33.8% increase over the worst-performing RF model (Dunn test p = 2.62 * 10^-10^, Cohen’s d = 11.637). At MAF 0.01, the GB model had a 14.3% increased accuracy than the second-best RF model (Dunn test p = 0.38, Cohen’s d = 6.48) and a 32.4% increase over the worst-performing GBLUP model (Dunn test p = 2.71 * 10^-9^, Cohen’s d = 13.253). Finally, at MAF 0.005, the GB model had a 14.8% increased accuracy than the second-best RF model (Dunn test p = 0.33, Cohen’s d = 5.758) and a 31.4% increase over the worst-performing GBLUP model (Dunn test p = 1.53 * 10^-9^, Cohen’s d = 12.153).

**Figure 1:**
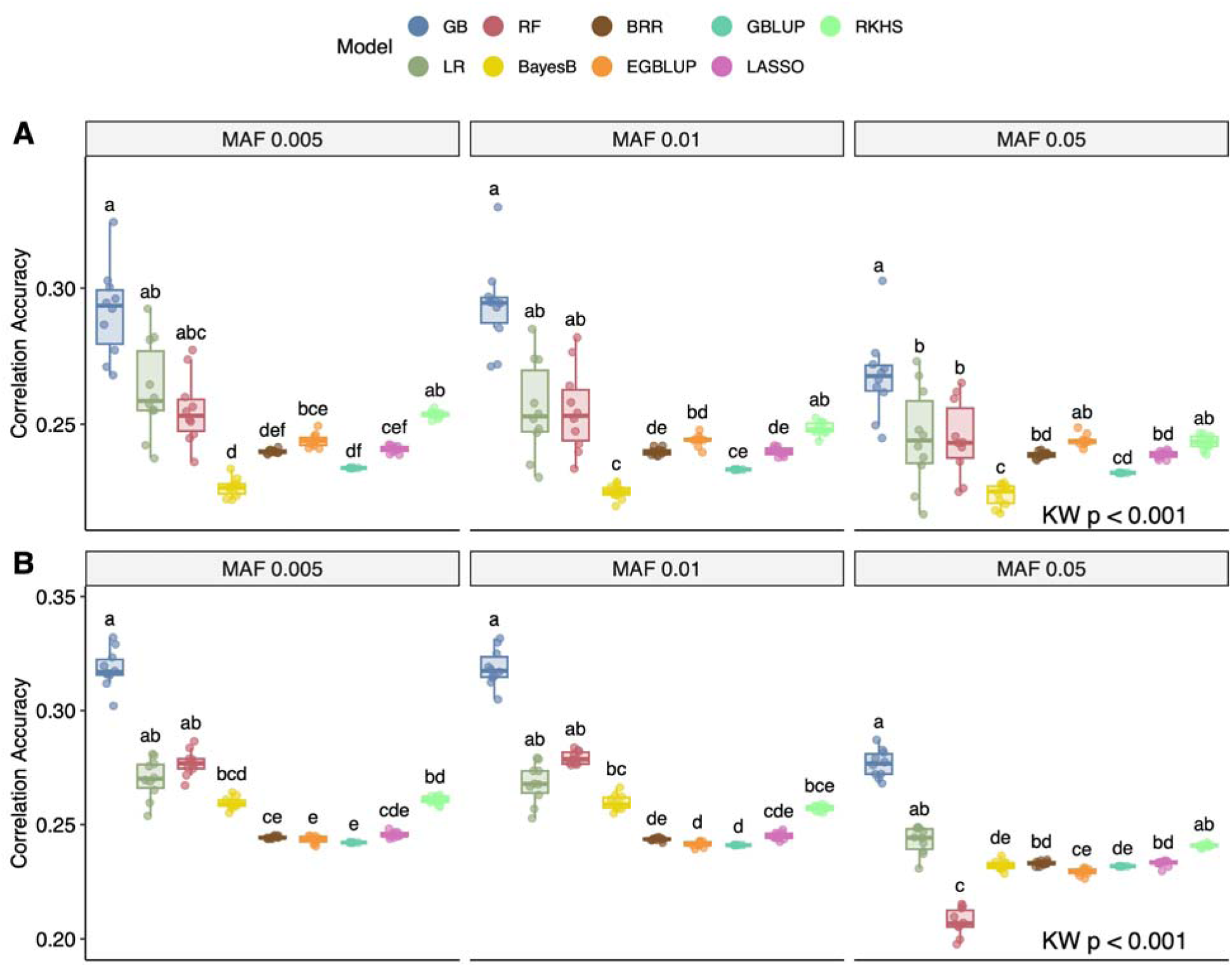
Differences in correlation accuracy among genomic selection models using MAF thresholds of 0.005, 0.01, and 0.05. Results are shown for training datasets comprising **(A)** only the F generation and **(B)** the full dataset. KW = Kruskal-Wallis test, which showed significant differences between model predictions at each MAF. Letters indicate significant differences based on Dunn’s post-hoc test corrected for multiple testing using the Benjamini-Hochberg method.

At MAFs of 0.01 and 0.005, all models except EGBLUP saw significant increases in correlation accuracy ranging from 1.5% (BRR) to 13.3% (BayesB) when trained on the entire dataset as opposed to just the F2 generation (p < 0.001, Wilcoxon signed-rank test) (Supplementary Figure 1, Supplementary Table 2).

Notably, there were much larger increases in correlation accuracy for all models when MAF was reduced from 0.05 to 0.01, from 5.2% (LASSO) to 34.6% (RF) (Figure 2, Supplementary Table 2). The best-performing GB model saw a 15.2% increase in correlation accuracy from 0.277 to 0.319 when trained on MAF 0.01 as opposed to MAF 0.05 (p = 1.08 * 10^-5^, Wilcoxon signed-rank test). None of the ML models saw significant increases in performance when MAF was further lowered to 0.005 (Figure 2A). The conventional models EGBLUP, GBLUP, and RKHS were the only models whose performance further increased significantly when trained on MAF 0.005 instead of MAF 0.01, but this improvement was substantially smaller, with a maximum increase of 1.4% for the RKHS model (p = 4.87 * 10^-4^, Wilcoxon signed-rank test) (Figure 2B). These results indicate that GB is the best-performing model, and correlation accuracy of all models significantly increased by reducing MAF from 0.05 to 0.01. Using the combined dataset also increased correlation accuracy, although this increase was less pronounced than reducing MAF threshold.

**Figure 2:**
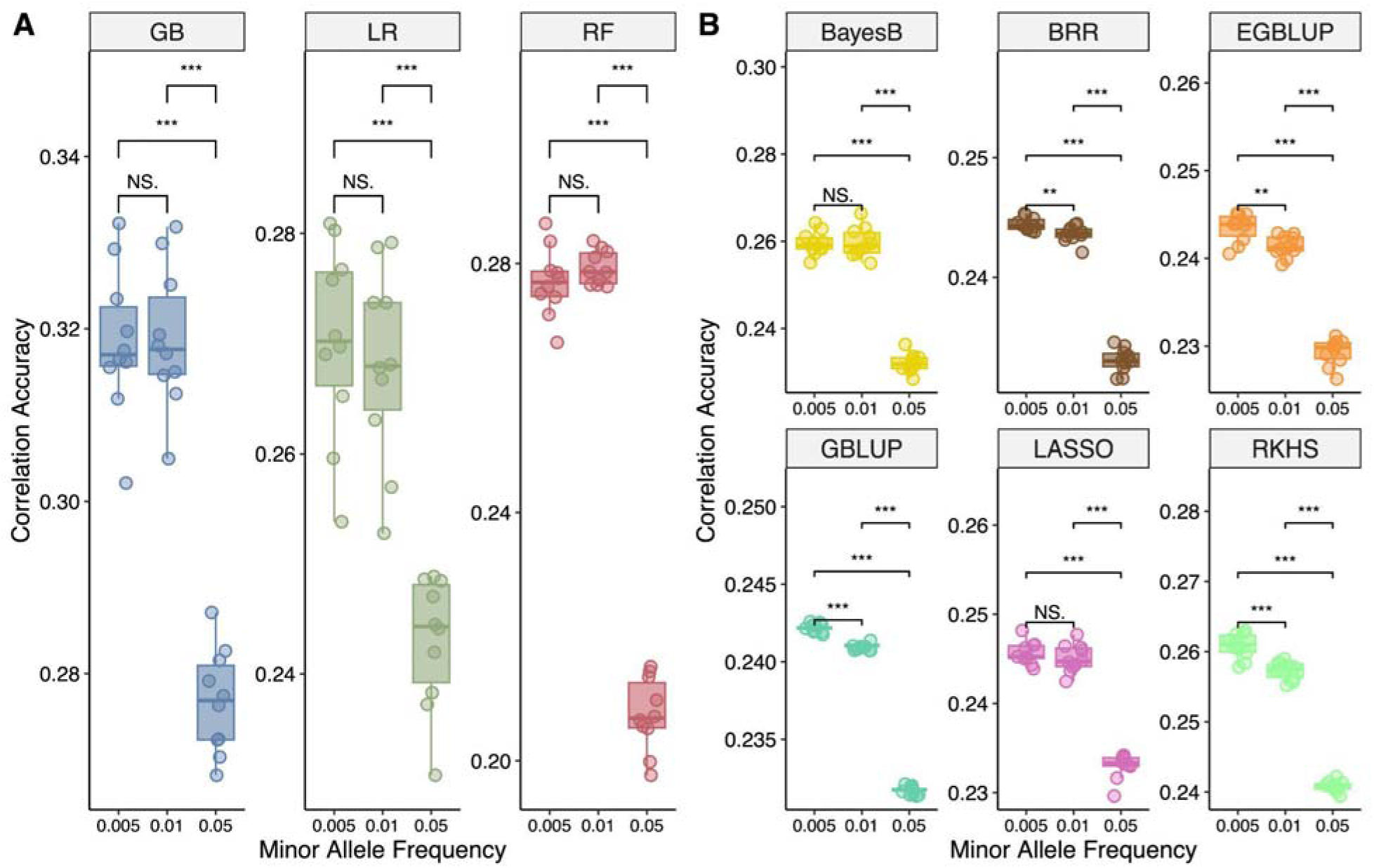
Box plots of correlation accuracy at MAF thresholds of 0.005, 0.01, and 0.05 for **(A)** machine learning models and **(B)** conventional genomic selection models. Significance levels are based on the Wilcoxon signed-rank test (*p < 0.05, **p < 0.01, ***p < 0.001; NS, not significant).

### ML model performance following hyperparameter tuning

We evaluated the performance of ML models when model hyperparameters were assigned default values rather than our standard approach using Bayesian optimization. The correlation accuracy of the best-performing GB model at MAF 0.01 increased 9.6% from 0.291 to 0.319 with hyperparameter tuning (p = 1.95 × 10^-3^, Wilcoxon signed-rank test), with correlation accuracy of all ML models significantly increasing between 8.5% and 25.1% when hyperparameters were tuned (all p < 0.001, Wilcoxon signed-rank test) (Supplementary Figure 2).

### Model performance following GSM addition

Given that including rare variants improved model performance, we evaluated whether targeted inclusion of strong-effect loci could further improve correlation accuracy. To do this, markers of strong effect identified by GWAS were added to the training dataset, resulting in the addition of 24 SNPs to the training population at MAF 0.05 and 31 SNPs added at MAF 0.01 and 0.005 (total n = 39,947, 48,174, 50,310 SNPs for n = 2,357, 2,363, and 2,363 individuals, respectively) [28]. These GSMs were missing from the initial training dataset either due to lower MAF or because their call rates were slightly below the QC threshold, but they all showed strong association with dermo resistance. As above, significant differences in model performances were observed at all three MAF filter levels following addition of GSMs: all-MAF 0.05 (Kruskal-Wallis p = 1.2 * 10^-15^); all-MAF 0.01 (Kruskal-Wallis p = 2 * 10-15); all - MAF 0.005 (Kruskal-Wallis p = 1.34 * 10^-15^) (Supplementary Figure 3). All models saw significant increases (p < 0.001, Wilcoxon signed-rank test) in correlation accuracy following the inclusion of GSMs (Figure 3, Supplementary Table 3).

**Figure 3:**
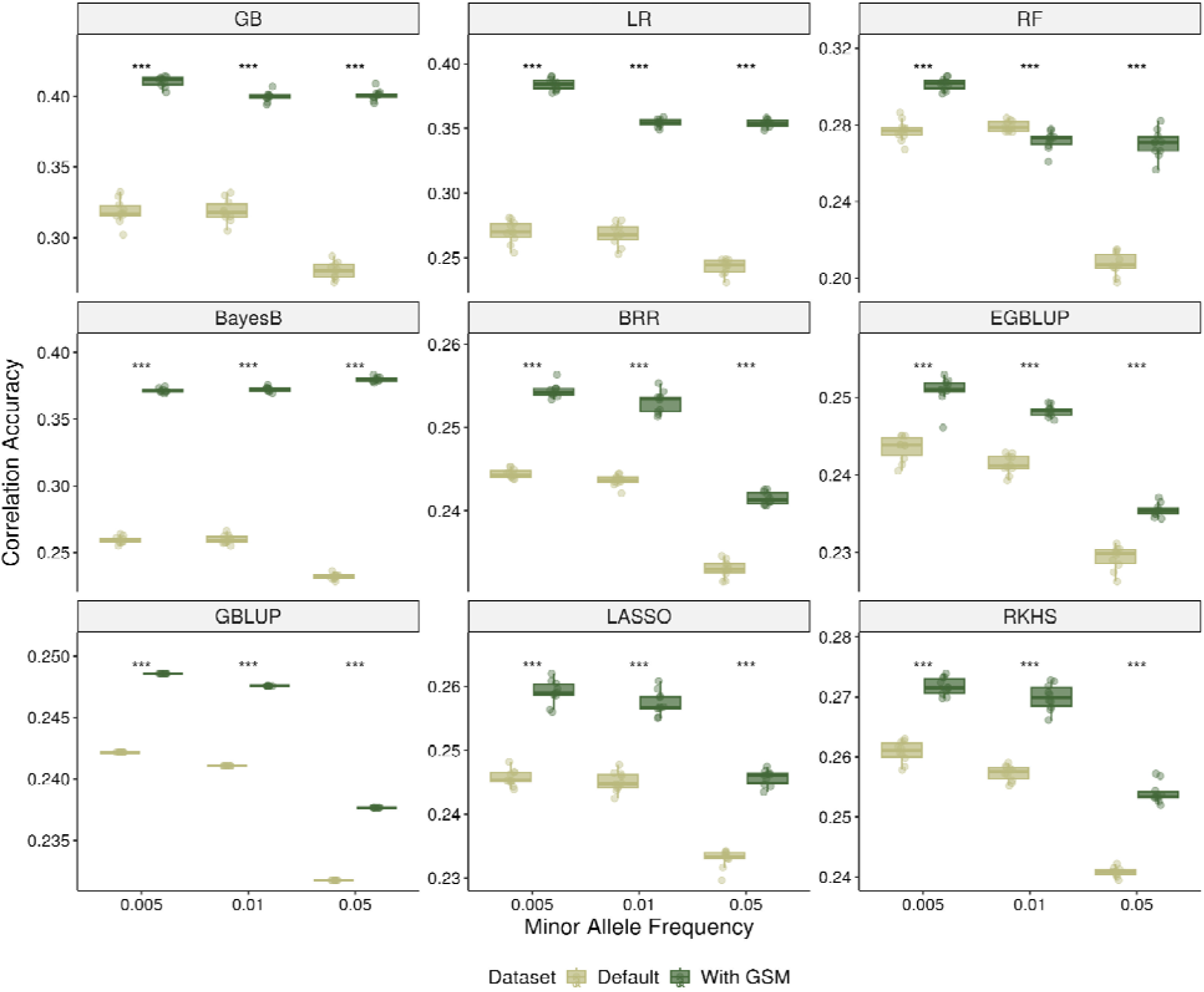
Box plots of correlation accuracy for each model trained on the default post-QC dataset versus a dataset including GWAS-selected markers (GSMs). Significance levels are based on the Wilcoxon signed-rank test (*p < 0.05, **p < 0.01, ***p < 0.001; NS, not significant).

The greatest increase in correlation accuracy at MAF 0.05 following addition of GSMs was realized by the GB (44.9% increase, p = 1.08 * 10^-5^, Wilcoxon signed-rank test), LR (45.7% increase, p = 1.8 * 10^-5^), RF (30.2% increase, p = 1.8 * 10^-5^) and BayesB models (63.7% increase, p = 1.8 * 10^-5^). The remaining conventional GS models modestly increased in correlation accuracy between 2.6% and 5.5%. GB models had the highest correlation accuracy of all models trained after addition of GSMs of 0.401 at MAF 0.05, 0.400 at MAF 0.01, and 0.410 at MAF 0.005, representing an additional increase in correlation accuracy of 2.5% from MAF 0.05 to MAF 0.005 (Supplementary Figure 3B). The correlation accuracy of the GB model at MAF 0.005 of 0.410 was 10.5% higher than the second-best BayesB model with a correlation accuracy of 0.371 (Dunn test p = 6.77 * 10^-10^, Cohen’s d = 13.351), and 63.4% higher than the worst-performing EGBLUP model with a correlation accuracy of 0.251 (Dunn test p = 4.72 * 10^-3^, Cohen’s d = 53.647). Overall, the best-performing GB model with GSMs at MAF 0.005 represents a 49.6% increase in correlation accuracy compared to the previous best single-generation GBLUP model in the F0 generation of 0.274 [19].

### Changes in GEBV predictions between models

Prior analyses of the F0 generation demonstrated strong agreement among five previously tested conventional GS models, with pairwise correlations between GEBV predictions all exceeding 0.88 [19]. In this study, the four new models evaluated (BayesB, GB, LR, RF) exhibited reduced correlations with those conventional GS models, with the lowest pairwise correlation between GB and EGBLUP at MAF 0.05 (Pearson ρ = 0.72) (Supplementary Figure 4A). Further, we observed notable changes in inter-model GEBV correlations following the addition of GSMs relative to previous experiments. The GB, LR, RF, and BayesB models with the largest improvements in correlation accuracy exhibited reduced GEBV correlations with the remaining conventional GS models, with the lowest observed correlation at MAF 0.005 occurring between the second-most accurate LR and least accurate EGBLUP models (Pearson ρ = 0.39) (Supplementary Figure 4B).

The GB, LR, RF, and BayesB models also showed significant shifts in their GEBV distributions following GSM addition consistent with improved discrimination between phenotypic classes. Visual inspection of the distributions revealed the emergence of secondary density peaks at lower GEBV values, corresponding to individuals classified as dead oysters, suggesting that these models more effectively captured genetic signals associated with mortality (Figure 4, Supplementary Figure 5). These changes reflected a restructuring of the underlying distribution of breeding values, with two-sample Kolmogorov– Smirnov tests confirming significant distributional changes for these models at all MAF thresholds (at MAF 0.005: GB p = 7.37 * 10^-26^, LR p = 5.71 * 10^-91^, RF p = 1.71 * 10^-3^, BayesB p = 1.87 *10^-27^). In contrast, no other models exhibited statistically significant changes in their GEBV distributions following GSM addition at any MAF thresholds (all p > 0.05, two-sample Kolmogorov–Smirnov test), indicating that distributional shifts were specific to models that most effectively leveraged additional GSMs (Figure 4, Supplementary Figures 6-7).

**Figure 4:**
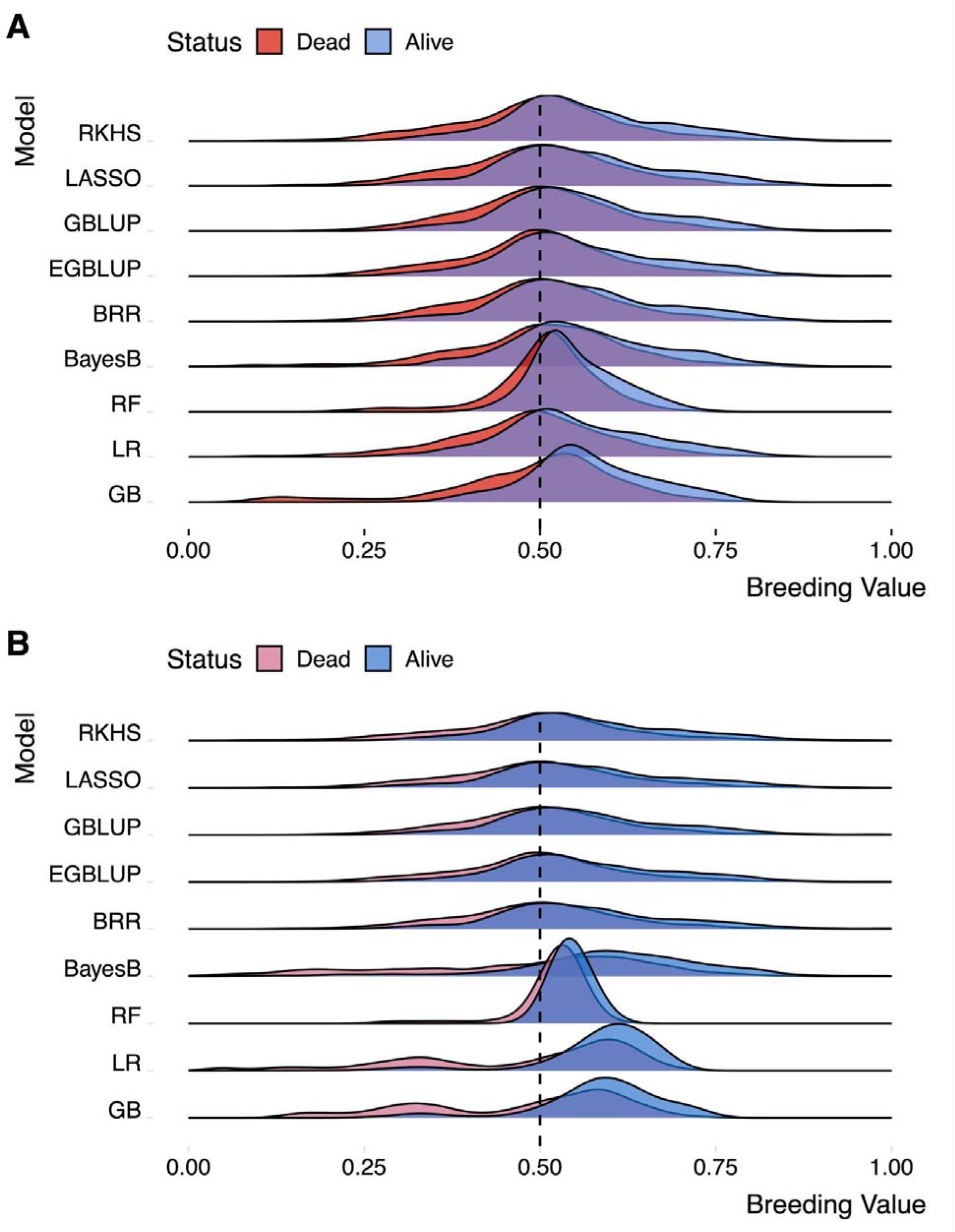
Distribution of genome-estimated breeding values (GEBVs) **(A)** before and **(B)** after the addition of GSMs to the training dataset at MAF = 0.005. The black dashed line indicates a breeding value of 0.50.

## Discussion

Dermo disease caused by the protozoan *Perkinsus marinus* remains a major constraint on oyster aquaculture production with the limited efficiency of traditional phenotypic selective breeding methods. Recently, we found that GS offers a promising alternative method that can improve the survival of oysters exposed to dermo more effectively than by phenotypic selection [19]. This study extends those findings to multiple generations and evaluates the potential of ML for genomic predictions on a pooled dataset from training populations of three successive generations. Here, we demonstrate that choices of dataset composition and modeling approach substantially influence performance, with GB models, lower MAF datasets, and GSMs all increasing correlation accuracy.

Increasing training dataset size is expected to improve model generalization performance by providing a more comprehensive representation of underlying variation in the real data distribution [39]. In this study, pooling individuals across three generations increased correlation accuracy for some models; however, at an MAF threshold of 0.05, most conventional GS models performed better when trained on the F2 generation alone, suggesting sensitivity to heterogeneity introduced by multiple generations of genomic data. In contrast, GB and BayesB models achieved higher correlation accuracy when trained on the full dataset. This pattern suggests these models were better able to capture generalizable signals of dermo resistance across generations, rather than overfitting to generation-specific patterns. Overall, reduced correlations between the newly evaluated models (BayesB, GB, LR, RF) and the five conventional GS models relative to the F0 generation indicate divergence in prediction patterns across models. Whereas previously tested models produced highly concordant GEBV rankings [19], the new models evaluated in this study generated distinct predictions and likely captured different aspects of underlying genetic architectures of dermo resistance.

In our previous single-generation study, model performance plateaued after inclusion of additional markers beyond the top 10K GWAS-identified SNPs [19], indicating that increasing marker density alone has limited impact on predictive accuracy if biologically informative variants are not retained. The present study shows that selective inclusion of rare variants through relaxed MAF thresholds significantly improves correlation accuracy. The larger sample size of the combined dataset (n = 2,357) provided greater statistical power to accurately identify rare variants, making it more defensible to reduce MAF QC threshold [28]. Lowering the MAF threshold from 0.05 to 0.01 introduced 8,220 additional SNPs and six individuals into the training dataset and resulted in large performance gains for all models except EGBLUP. The comparatively smaller gains observed when further lowering MAF from 0.01 to 0.005 reflect diminishing returns as fewer informative variants are introduced. These results reinforce that optimal QC thresholds are context-dependent and should be tailored to dataset characteristics. The large increase in correlation accuracy from MAF 0.05 to 0.01 suggests an important role for loci with rare alleles in the genetic architecture of dermo resistance. One plausible explanation is that these minor alleles are deleterious, recessive variants that disproportionately affect fitness, including resistance to dermo infection. Such variants have been extensively characterized in human disease-related traits [40, 41, 42], and may contribute to oyster mortality under dermo infection. Minor allele frequencies were higher in the dead population than in survivors for all markers with strong associations to dermo resistance, as expected for deleterious lethal alleles. Similarly, genotype distributions for these markers followed expected patterns of deleterious recessive effects, with heterozygous individuals showing reduced survival and homozygous minor genotypes being largely absent among survivors [28]. This result and hypothesis are consistent with the high levels of polymorphism or genetic load observed in oyster genomes [43, 44].

Our results also demonstrated clear increased correlation accuracy with inclusion of up to 31 GSMs, supporting previous findings that markers identified by GWAS can boost GS accuracy [29, 30, 31]. While dermo resistance is highly polygenic, increased statistical power from the multigenerational dataset enabled detection of these GSMs with larger effects. One of the GSMs that was initially excluded because of a slightly lower call rate, for example, accounted for about 8.1% of the phenotypic variation [28].

Consequently, more variation was accounted for upon GSM inclusion, creating a genetic architecture well suited to models that accommodate sparse or heterogeneous effect sizes. This is consistent with strong performance gains following GSM addition observed for BayesB, which can shrink weights of small-effect SNPs to zero [45, 46], and GB, which flexibly captures complex interactive effects within the underlying dataset [47]. Our results also revealed that GSM addition further altered the underlying ranking structure of breeding values predicted by high-performing models (GB, LR, RF, and BayesB).

Notably, the incorporation of GSMs further decreased inter-model GEBV correlations between these and other models to as low as 0.39 at MAF 0.005. The observed changes in GEBV distributions for these models following GSM addition suggest that they produced more confident predictions indicative of a shift from approximately normal distributions of breeding values toward more polarized distributions, reflecting clearer separation between classes for a binary prediction task.

GB models achieved the highest overall correlation accuracy, outperforming alternative approaches by at minimum 10.5% across all training scenarios. These results are consistent with prior studies reporting increased effectiveness of GB frameworks for various genomic prediction tasks [48, 49, 50]. GB fuses multiple weak learners into a strong ensemble over many iterations to learn complex nonlinear interactions present within data structures and performs well on tabular data [51]. The highest correlation accuracy in this study was achieved using GB on the most comprehensive dataset, incorporating all three generations, the lowest MAF threshold, and additional GSMs. This indicates that GB is capable of leveraging increased feature space without substantial overfitting by prioritizing informative markers while down-weighting noise. The implementation of GB used in this study, XGBoost, is computationally efficient and scalable, provides built-in regularization techniques against overfitting, and can be used for both classification and regression tasks [52]. Altogether, our results strengthen the motivation for future tests of GB and other ML models in genomic predictions for aquaculture applications.

The strong performance of ML models observed here also underscores the importance of appropriate hyperparameter tuning in realizing genomic prediction gains with ML [53]. For example, GB requires careful optimization of hyperparameters such as learning rate, number of iterations, and regularization terms, all of which can substantially influence model accuracy and are difficult to determine *a priori* without empirical tests [54]. Accordingly, in this study, all ML models exhibited substantial improvements (8.5–25.1%) in correlation accuracy following hyperparameter tuning via Bayesian optimization. Bayesian optimization balances exploration of large hyperparameter spaces with exploitation of promising hyperparameter combinations and thereby has performance and efficiency advantages over traditional grid or random hyperparameter search methods in genomic prediction applications [55].

An important avenue for future research beyond the scope of this study is the integration of ML approaches with variant discovery frameworks. Interpretable or “white box” ML models, such as GB, LR, and RF, can provide measures of feature importance that offer biological insight into model predictions, which may complement or extend traditional genome scans or GWAS by identifying outlier loci associated with phenotypic variation [56, 57, 58]. Several ML–based feature selection methods exist that prioritize informative subsets of markers and may help address the “large p, small n” challenges inherent in high-dimensional genomic datasets [59, 60]. Emerging results suggest such approaches can outperform conventional GWAS-based feature selection [32, 61, 62]. Future studies in aquaculture can seize the opportunity to integrate interpretable ML methods with genomic analyses, enabling more reliable identification of key genetic markers while improving prediction performance in limited-sample training datasets.

Altogether, our results suggest context dependence for model performances and the importance of considering the underlying trait architecture. The increase in correlation accuracy from the previous peak of 0.274 to 0.410 in this study represents a 49.6% increase for future applications of GS for dermo resistance in Eastern oysters. This result is promising for selective breeding efforts, considering the previous-best F0 generation GBLUP model already produced increased survival against lab-based dermo challenge in F1 progeny compared to phenotypic selection and a genomically down-selected control [19]. Field evaluations of genomically selected progeny in this study are ongoing and will inform how model predictions translate to real-world scenarios and stressors. Improved performance in this study using ML models emphasizes that continued tests of ML-based approaches for genomic prediction are needed, with considerable rigor applied to hyperparameter tuning and cross-validation comparisons with existing approaches.

## Materials and Methods

### Sample collection, data generation, and QC

In 2022, wild Eastern oysters were collected from Florida and challenged with dermo to create a training population for genomic selection [19]. Briefly, dermo challenge was conducted in the lab by injecting *P. marinus* into the shell cavity. The challenge experiment continued until ∼50% mortality was reached, upon which phenotypes (dead or live) were recorded. Oysters from both the training population and the unchallenged Florida wild breeding population were genotyped with the 66K OysterCv SNP array [24]. The training population was used to evaluate the different datasets and models, and the best approach was used to predict genomic estimated breeding values (GEBVs) of the breeding population. Oysters with the highest GEBVs were selected to produce the genomic selected F1 group along with control groups [19]. This process was repeated for the F1 and F2 generations, creating three training populations and three generations of genomic selected and control groups [28]. The current study collected and analyzed the training datasets of all three generations, F0, F1, and F2. Across three training populations, we genotyped a total of 2423 oysters at 65893 SNPs, with a total of 505 samples from the F0 generation, 940 samples from the F1 generation, and 978 samples from the F2 generation, respectively. We removed training samples with an individual call rate < 90%, defined as successful genotype calls at below 90% of assayed SNP loci. We further removed any SNP markers with a < 95% call rate and MAF < 0.05. Any remaining missing genotypes were imputed using the mean genotype value at each SNP per generation.

### Identification of strong-effect markers using genome-wide association studies (GWAS)

In another study, GWAS was performed on the samples from this dataset and identified markers significantly associated with dermo resistance after correcting for multiple comparisons at the chromosomal level [28]. Some of these markers were initially excluded during QC because of their slightly low call rate. Of these markers, 31 showed reasonable call rates (>85%) and their biological significance was validated through ontology, cross-generational trends, and field performance. We added these 31 GWAS-selected markers (GSMs) into the training dataset, of which 24 passed filtering at MAF 0.05 and all 31 were retained at MAF 0.01 and 0.005. We resolved any missing values for GSMs using nearest-neighbor imputation from the 10 individuals with highest genetic relatedness within each generation; similar methods relying on genetic proximities or parentages have been shown to produce greater genomic prediction accuracy than mean imputation [63].

### Genomic selection model training

We trained and evaluated a total of nine different GS models. These included Bayesian B (BayesB), Bayesian ridge regression (BRR) [64], genomic best linear unbiased prediction (GBLUP) [65], extended GBLUP (EGBLUP) [66], Bayesian least adjusted selection shrinkage operator (LASSO) [67], and reproducing kernel Hilbert space (RKHS) [68] models trained in R 4.4.0 using BWGS v0.2.0 [69] and BGLR v1.1.4 [70] packages. We trained gradient boosting (GB), logistic regression (LR), and random forest (RF) supervised machine learning (ML) models in Python 3.12.2 using scikit-learn 1.6.1 [71] and xgboost 3.0.4 [51]. Previously, we tested GBLUP, EGBLUP, LASSO, BRR, and RKHS on the F0 generation training population, with descriptions of these models’ underlying architectures and their respective performance available in [19].

Here, we also evaluated the effectiveness of four additional models, BayesB, GB, LR, and RF, for genomic predictions of dermo resistance. BayesB is a sparse Bayesian variable selection model which assumes a mixture prior in which a proportion of markers has zero effect and the rest follow a gamma or exponential distribution, enabling heterogeneous shrinkage of marker effects [72]. LR models the probability of a binary outcome by using a logit link function, estimating predictor effects through maximum likelihood [73]. RF is an ensemble learning method that combines multiple decision trees built on bootstrap samples with random feature selection to improve accuracy and reduce overfitting, also known as bootstrap aggregation or bagging [74]. GB is an iteration-based ensemble technique that sequentially fits weak decision trees to the residuals of prior models over time to build a strong learner and minimize prediction error, a technique known as boosting [75].

All models predicted genome-estimated breeding values (GEBVs) as a continuous variable between 0 and 1, functioning as a probability estimate of organismal survival against dermo infection. As is standard in binary genomic prediction tasks, model correlation accuracy was defined as the Pearson correlation between the GEBV and the true phenotype of either 0 (dead) or 1 (alive) at the conclusion of the dermo challenge. For conventional models trained in R, we split the training data into an 80% training set and 20% testing set. We ran 10 randomized iterations of stratified 5-fold cross-validation split into 80% training and 20% testing data for each fold and predicted GEBVs of the 20% unseen test fold.

For ML model training, we first split the training data into an 80% training set and 20% validation set for hyperparameter tuning. Hyperparameter tuning refers to the process of determining optimal parameters which influence model training, such as the number of training iterations. We tuned hyperparameters using 100 iterations of Bayesian optimization using scikit-optimize 0.10.2 aimed at optimizing model correlation accuracy, with each iteration undergoing five-fold cross-validation [76]. Bayesian optimization is a sequential model-based approach that uses a probabilistic surrogate function to model the objective surface and an acquisition function to efficiently select the next set of hyperparameters to evaluate, thereby balancing exploration of hyperparameter search space with exploitation of promising regions. Hyperparameter search spaces for all models trained in this study are listed in Supplementary Table 1, with final hyperparameter values available at (https://github.com/henrysun9074/gsAI/tree/main/MLmodels). After hyperparameter tuning, we reshuffled the data and again ran 10 randomized iterations of stratified 5-fold cross-validation with an 80-20 train-test split.

### Genomic prediction model evaluation

We compared correlation accuracy across models under several conditions: training on the full dataset (i.e., F0, F1, and F2) versus only the most recent F2 generation; at MAF thresholds of 0.05, 0.01, and 0.005; for ML models, with and without hyperparameter tuning; finally, for all models, with or without addition of GSMs. When evaluating ML models trained without hyperparameter tuning, default hyperparameter values from scikit-learn were used. Model performance was evaluated over 10 iterations, with correlation accuracy reported as the mean accuracy across iterations. Within-model comparisons of correlation accuracy were conducted using Wilcoxon signed-rank tests to evaluate differences between (i) training populations (F2 generation vs. full dataset), (ii) MAF thresholds, (iii) hyperparameter tuning methods (default versus Bayesian optimization) for ML models, and (iv) addition of GSMs. Between-model differences in correlation accuracy were assessed using the Kruskal–Wallis test. Post-hoc pairwise comparisons were performed using Dunn’s test adjusted for multiple comparisons using the Benjamini– Hochberg procedure to control the false discovery rate [77], with effect sizes quantified using Cohen’s d-statistic [78]. Nonparametric methods were used due to unequal variances as confirmed by Levene’s test for both prediction accuracies and GEBVs, with statistical significance assessed at α = 0.05. Agreement between GEBVs predicted by different models was assessed using Pearson correlation coefficients and two-sample Kolmogorov-Smirnov tests. All data analysis and visualization were performed in R 4.4.0 [79].

## Data Availability Statement

Genotypic and phenotypic data from challenged oysters are available upon request. All code for genomic selection model tuning, training, and cross-validation, and subsequent statistical analyses and figure generation from this study is available at https://github.com/henrysun9074/gsAI. Model weights saved as joblib files for trained machine learning models are also available upon request.

## Acknowledgements

We thank the Center for Aquaculture Technology for their service of DNA extraction and SNP array genotyping. This study was funded by the Defense Advanced Research Projects Agency (DARPA) under Contract No. HR0011122C0136 and partly by the National Oceanic and Atmospheric Administration (NOAA), United States Department of Commerce through the Atlantic States Marine Fisheries Commission (award NA18NMF4720321). XG, DB, and JL are also supported by USDA NIFA Hatch projects 1021665/NJ30401, 1009201/NJ32114, and LAB94509, respectively. HS is supported by the NSF Graduate Research Fellowship under grant DGE-2139754. We are grateful to Callie Hundley for providing helpful comments on the manuscript.

## Author Contributions

XG, DB, JLP, and SR designed experiments. SC and JLP conducted disease challenge experiments. SR and MLW produced and cultured experimental oysters. XG and ZW conducted genotyping. HS and PC trained genomic selection models and analyzed data. DB, XG, JLP, and SR acquired funding. XG and JW supervised the study. HS wrote the manuscript with inputs from PC, JW, and XG. All authors contributed to the review and editing of the manuscript and approved the final version for publication.

