## Supplementary Material for "Multigenerational machine learning-based genomic prediction for dermo resistance in eastern oyster *Crassostrea virginica*"

**Supplementary Table 1:** Hyperparameter search space for each of the three machine learning models tuned using Bayesian optimization. Integer (int) and continuous (real) parameters were sampled from within defined ranges, while categorical parameters were selected from specified discrete values. For GB models, the tree construction method was set to ‘hist’, and model performance was evaluated using the ‘logloss’ metric. For LR models, the maximum number of iterations was set to 1000, and the ‘saga’ solver was used.

| **Model** | **Hyperparameter** | **Search Space** |
| --- | --- | --- |
| LR | C | real(0.00001, 10, prior="log-uniform") |
|  | penalty | categorical([L1, L2]) |
| RF | n_estimators | int(100,2000) |
|  | max_depth | int(3,50) |
|  | max_features | categorical(["sqrt", "log2"]) |
|  | min_samples_split | int(2,20) |
|  | min_samples_leaf | int(1,10) |
| GB | n_estimators | int(100,2000) |
|  | max_depth | int(3,15) |
|  | learning_rate | real(0.001, 0.3) |
|  | subsample | real(0.5,1.0) |
|  | colsample_bytree | real(0.5,1.0) |
|  | min_child_weight | int(1,10) |
|  | gamma | real(0,5) |

**Supplementary Table 2:** Average correlation accuracies across 10 iterations of 5-fold cross validation for all genomic selection models trained on both the most recent F2 generation and all generations, as well as different MAF filters of 0.05, 0.01, and 0.005.

| **MAF** | **Model** | **Generation** | **Average Correlation Accuracy** |
| --- | --- | --- | --- |
| 0.005 | GB | F2 | 0.291 |
| 0.005 | GB | all | 0.318 |
| 0.005 | LR | F2 | 0.263 |
| 0.005 | LR | all | 0.270 |
| 0.005 | RF | F2 | 0.255 |
| 0.005 | RF | all | 0.277 |
| 0.005 | BayesB | F2 | 0.227 |
| 0.005 | BayesB | all | 0.260 |
| 0.005 | BRR | F2 | 0.24 |
| 0.005 | BRR | all | 0.244 |
| 0.005 | EGBLUP | F2 | 0.244 |
| 0.005 | EGBLUP | all | 0.243 |
| 0.005 | GBLUP | F2 | 0.234 |
| 0.005 | GBLUP | all | 0.242 |
| 0.005 | LASSO | F2 | 0.241 |
| 0.005 | LASSO | all | 0.246 |
| 0.005 | RKHS | F2 | 0.254 |
| 0.005 | RKHS | all | 0.261 |
| 0.01 | GB | F2 | 0.294 |
| 0.01 | GB | all | 0.319 |
| 0.01 | LR | F2 | 0.256 |
| 0.01 | LR | all | 0.268 |
| 0.01 | RF | F2 | 0.255 |
| 0.01 | RF | all | 0.279 |
| 0.01 | BayesB | F2 | 0.225 |
| 0.01 | BayesB | all | 0.260 |
| 0.01 | BRR | F2 | 0.240 |
| 0.01 | BRR | all | 0.244 |
| 0.01 | EGBLUP | F2 | 0.244 |
| 0.01 | EGBLUP | all | 0.241 |
| 0.01 | GBLUP | F2 | 0.233 |
| 0.01 | GBLUP | all | 0.241 |
| 0.01 | LASSO | F2 | 0.240 |
| 0.01 | LASSO | all | 0.245 |
| 0.01 | RKHS | F2 | 0.248 |
| 0.01 | RKHS | all | 0.257 |
| 0.05 | GB | F2 | 0.268 |
| 0.05 | GB | all | 0.277 |
| 0.05 | LR | F2 | 0.245 |
| 0.05 | LR | all | 0.243 |
| 0.05 | RF | F2 | 0.245 |
| 0.05 | RF | all | 0.207 |
| 0.05 | BayesB | F2 | 0.224 |
| 0.05 | BayesB | all | 0.232 |
| 0.05 | BRR | F2 | 0.239 |
| 0.05 | BRR | all | 0.233 |
| 0.05 | EGBLUP | F2 | 0.244 |
| 0.05 | EGBLUP | all | 0.229 |
| 0.05 | GBLUP | F2 | 0.232 |
| 0.05 | GBLUP | all | 0.232 |
| 0.05 | LASSO | F2 | 0.239 |
| 0.05 | LASSO | all | 0.233 |
| 0.05 | RKHS | F2 | 0.243 |
| 0.05 | RKHS | all | 0.241 |

**Supplementary Table 3:** Average correlation accuracies across 10 iterations of 5-fold cross validation for all genomic selection models trained with datasets including GSMs at different MAF filters; all models were trained on all three generations of data.

| **MAF** | **Model** | **Average Correlation Accuracy** |
| --- | --- | --- |
| 0.005 | GB | 0.410 |
| 0.005 | LR | 0.384 |
| 0.005 | RF | 0.301 |
| 0.005 | BayesB | 0.371 |
| 0.005 | BRR | 0.254 |
| 0.005 | EGBLUP | 0.251 |
| 0.005 | GBLUP | 0.251 |
| 0.005 | LASSO | 0.259 |
| 0.005 | RKHS | 0.272 |
| 0.01 | GB | 0.400 |
| 0.01 | LR | 0.354 |
| 0.01 | RF | 0.272 |
| 0.01 | BayesB | 0.372 |
| 0.01 | BRR | 0.253 |
| 0.01 | EGBLUP | 0.248 |
| 0.01 | GBLUP | 0.248 |
| 0.01 | LASSO | 0.257 |
| 0.01 | RKHS | 0.270 |
| 0.05 | GB | 0.401 |
| 0.05 | LR | 0.354 |
| 0.05 | RF | 0.270 |
| 0.05 | BayesB | 0.380 |
| 0.05 | BRR | 0.241 |
| 0.05 | EGBLUP | 0.235 |
| 0.05 | GBLUP | 0.238 |
| 0.05 | LASSO | 0.246 |
| 0.05 | RKHS | 0.254 |

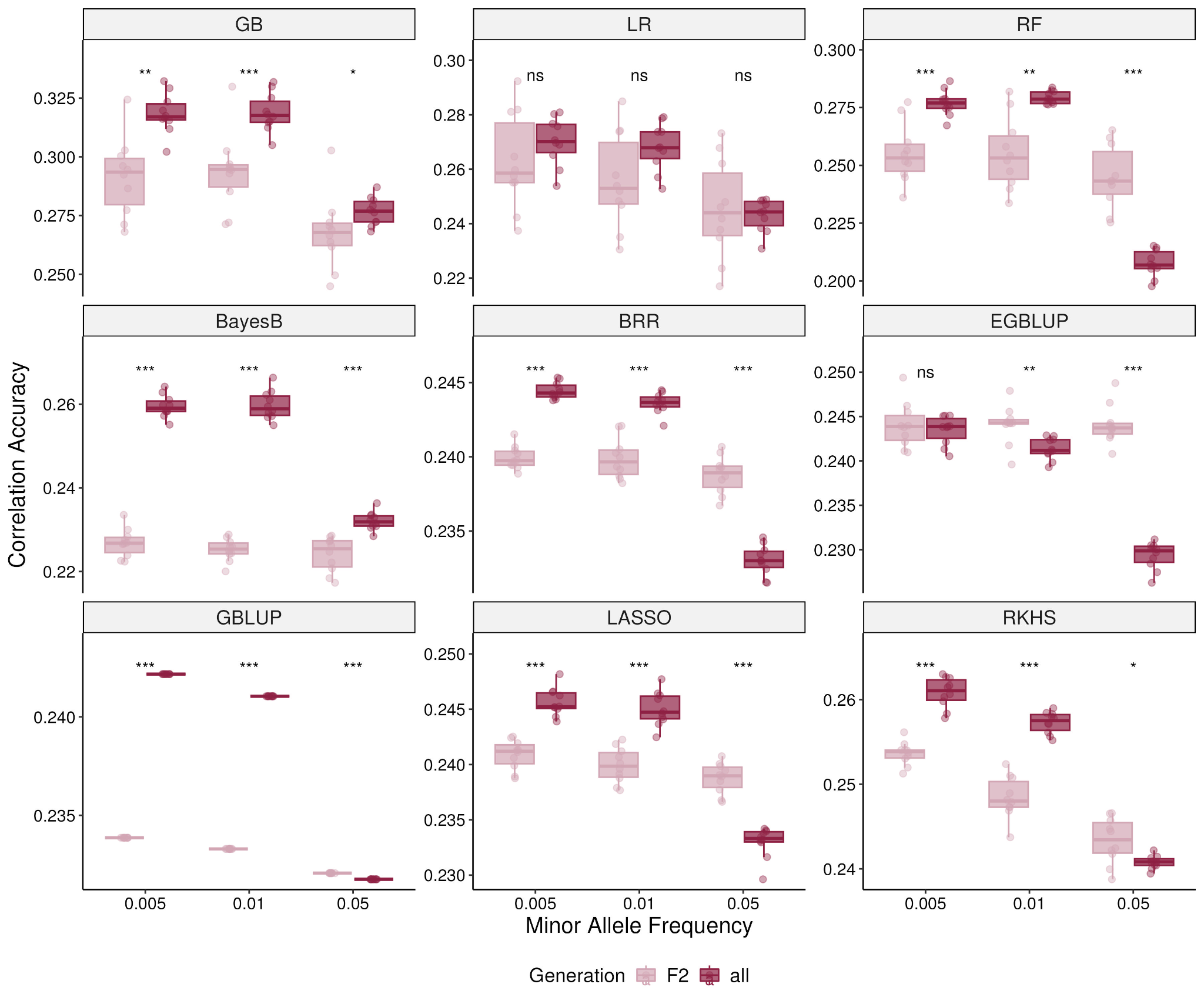
**Supplementary Figure 1:** Box plots of model correlation accuracy across all MAF thresholds comparing training on the F₂ generation versus the full dataset. Significance levels are based on the Wilcoxon signed-rank test (*p < 0.05, **p < 0.01, ***p < 0.001; NS, not significant).

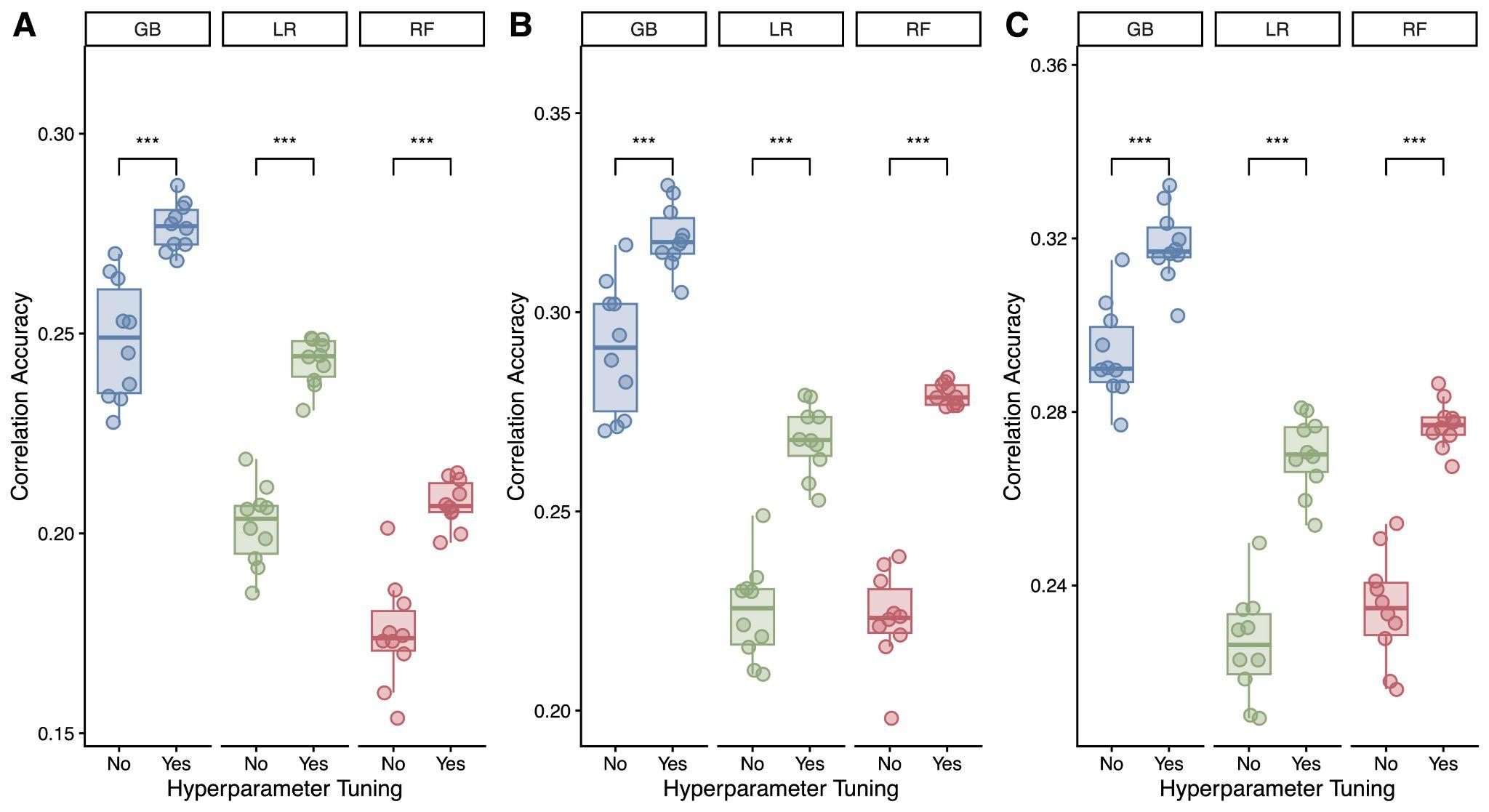

**Supplementary Figure 2:** Effect of hyperparameter tuning conducted using 100 iterations of Bayesian optimization on correlation accuracy at MAF thresholds of **(A)** 0.05, **(B)** 0.01, and **(C)** 0.005. Models trained without hyperparameter tuning used default values from the scikit-learn package. See Supplementary Table 1 for the list of hyperparameters optimized for each model. Significance levels are based on the Wilcoxon signed-rank test (*p < 0.05, **p < 0.01, ***p < 0.001; NS, not significant).

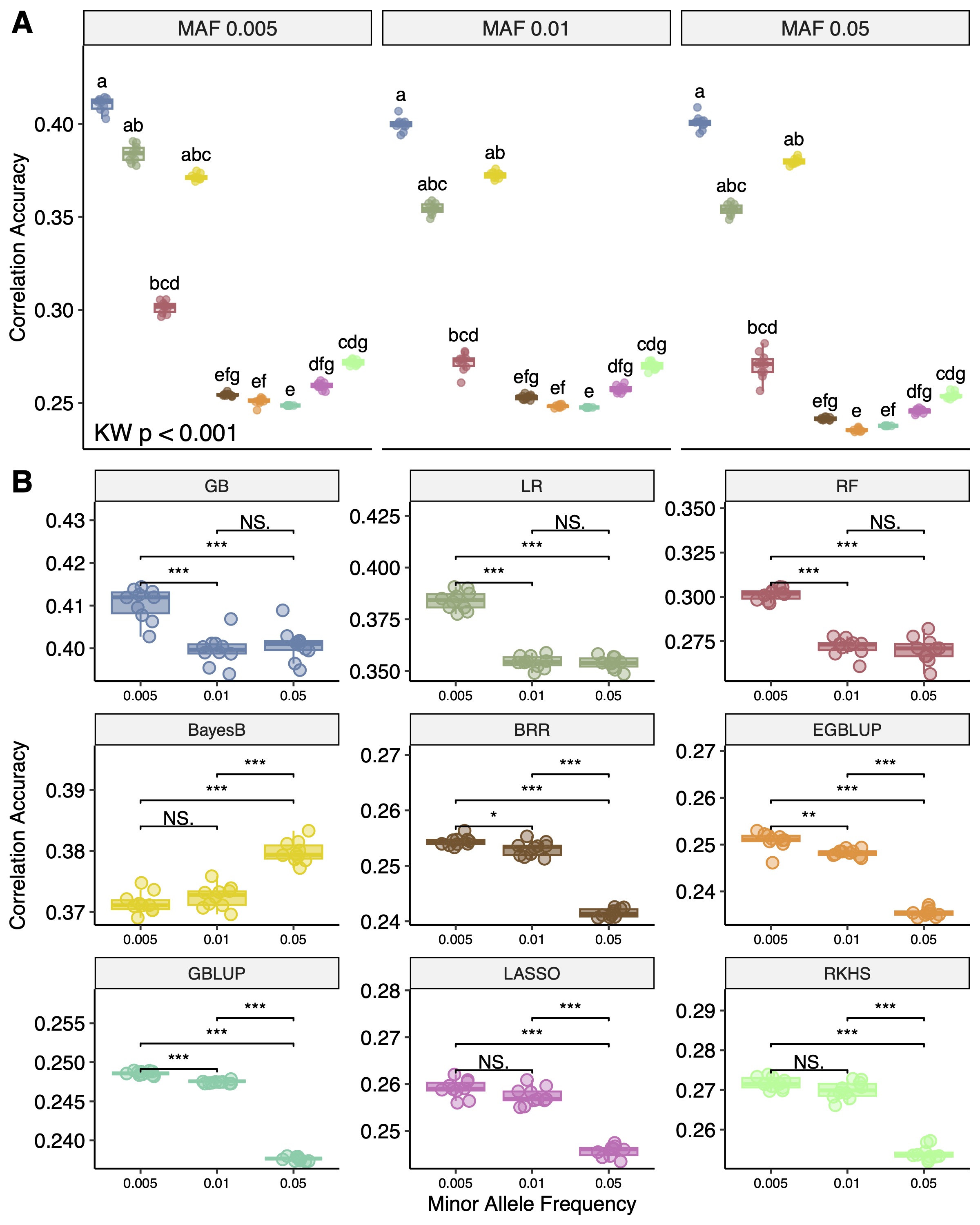

**Supplementary Figure 3: (A)** Differences in correlation accuracy among genomic selection models following addition of GWAS-selected markers (GSMs) using minor allele frequency (MAF) thresholds of 0.005, 0.01, and 0.05. KW = Kruskal-Wallis test, which showed significant differences between model predictions at each MAF. Letters indicate significant differences based on Dunn’s test corrected for multiple testing using the Benjamini-Hochberg method**. (B)** Box plots of model performance with addition of GSMs at each MAF. Significance levels are based on the Wilcoxon signed-rank test (*p < 0.05, **p < 0.01, ***p < 0.001; NS, not significant).

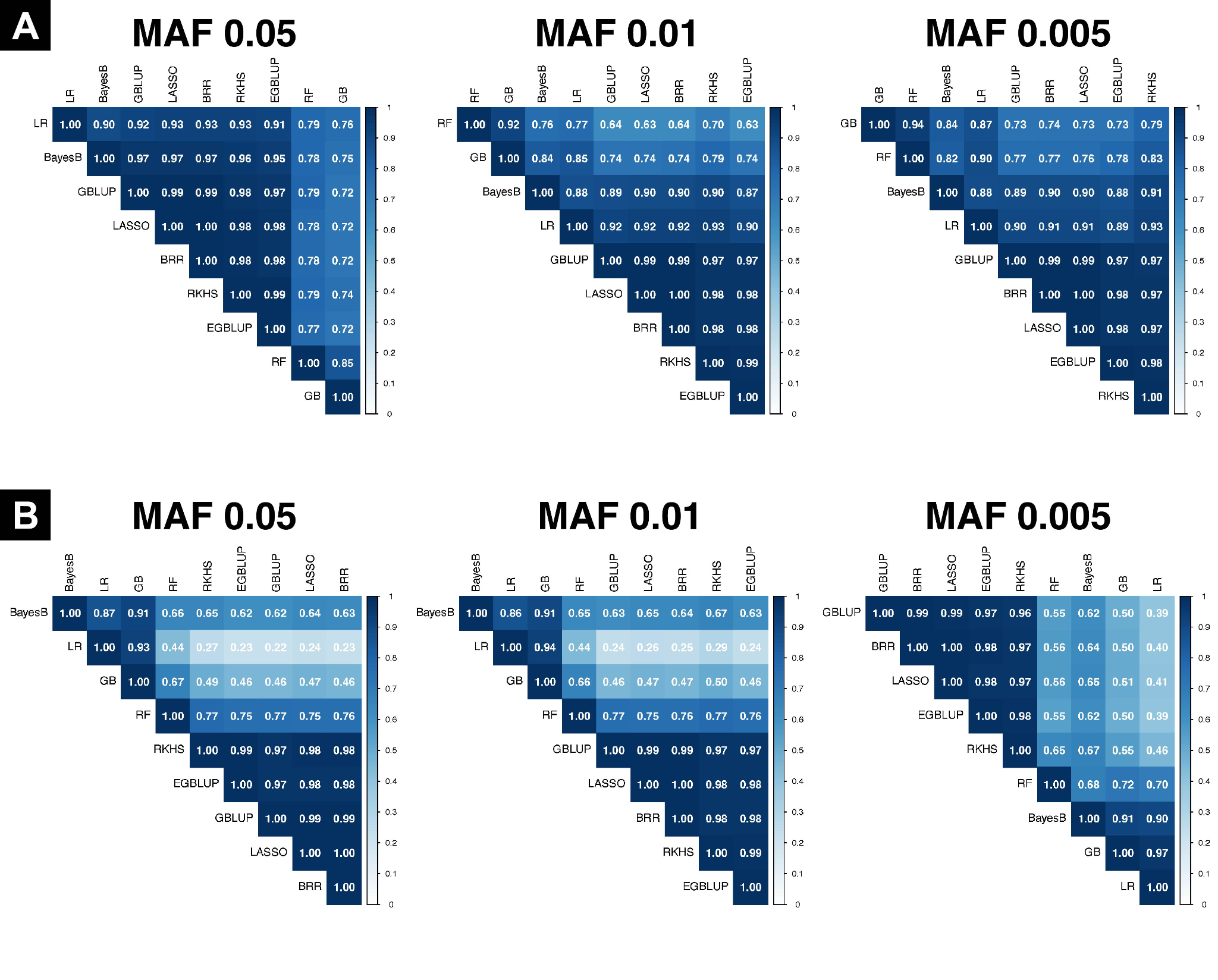

**Supplementary Figure 4:** Pairwise Pearson correlations of genomic estimated breeding values (GEBVs) at each MAF threshold for **(A)** the default dataset and **(B)** the dataset including GSMs.

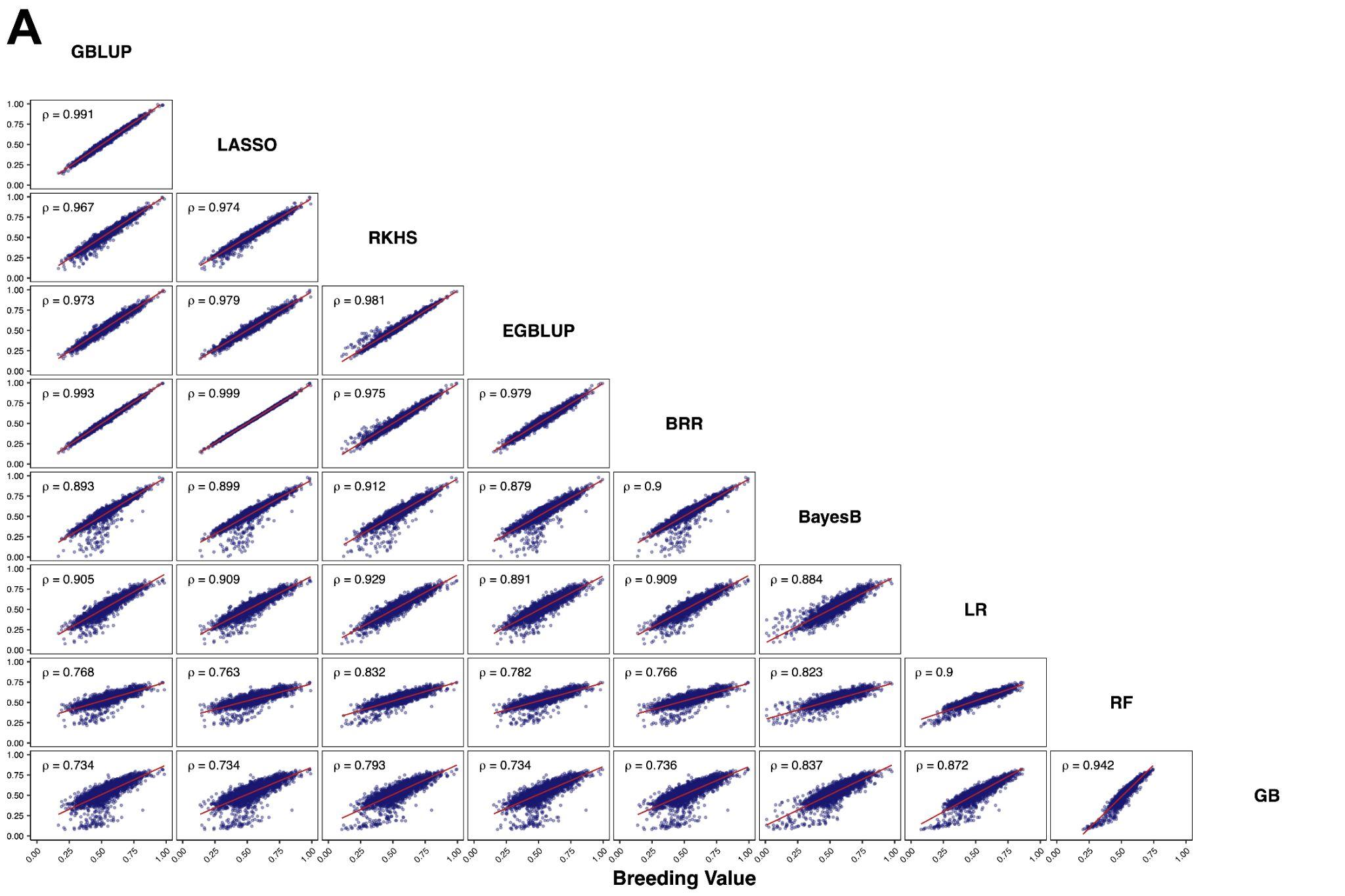

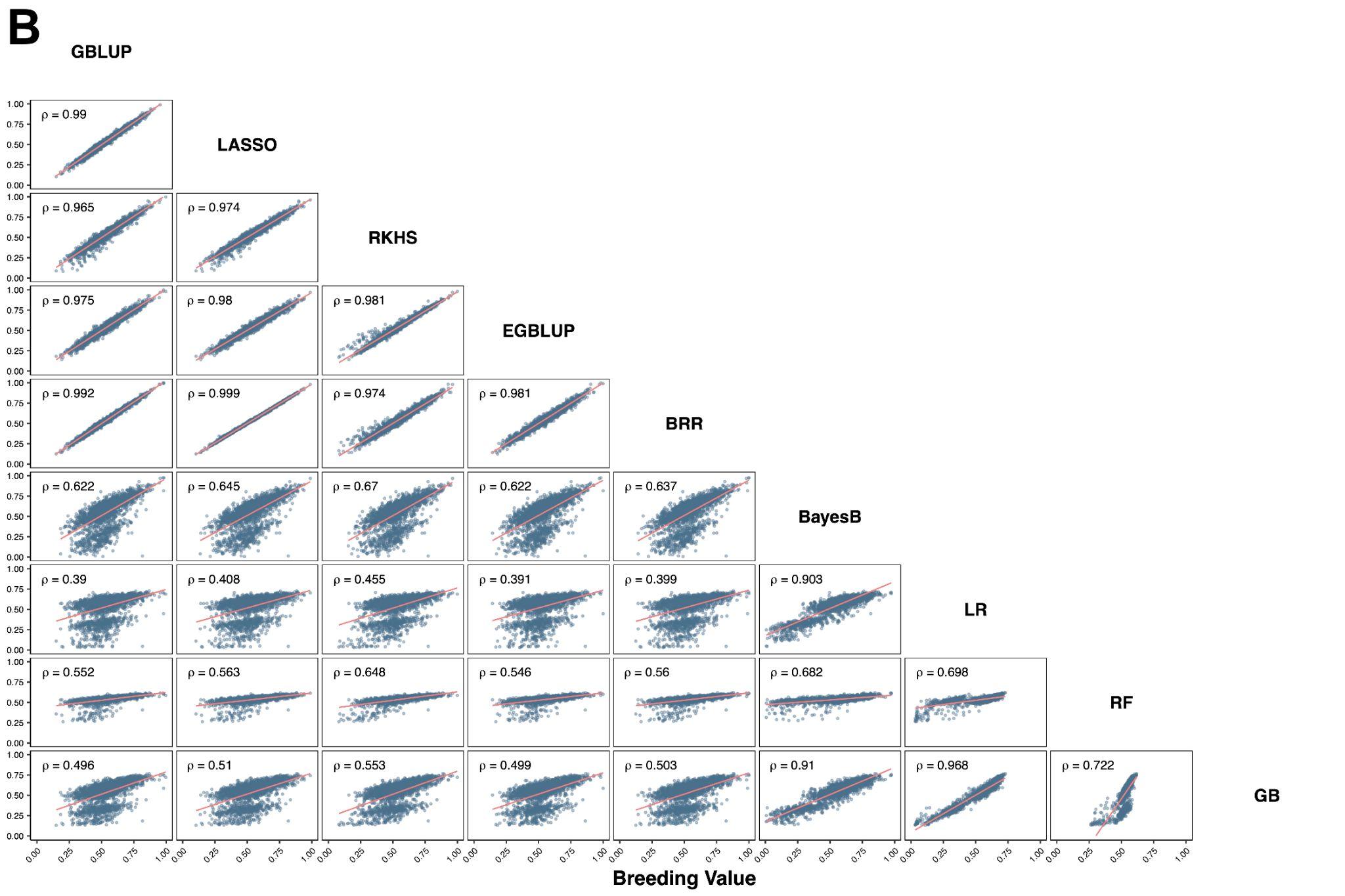

**Supplementary Figure 5:** Scatterplot matrix (SPLOM) of GEBVs at MAF = 0.005 for **(A)** the default dataset **(B)** with GSMs included. Inset values on each scatterplot indicate Pearson correlation between model breeding value predictions for each pairwise comparison.

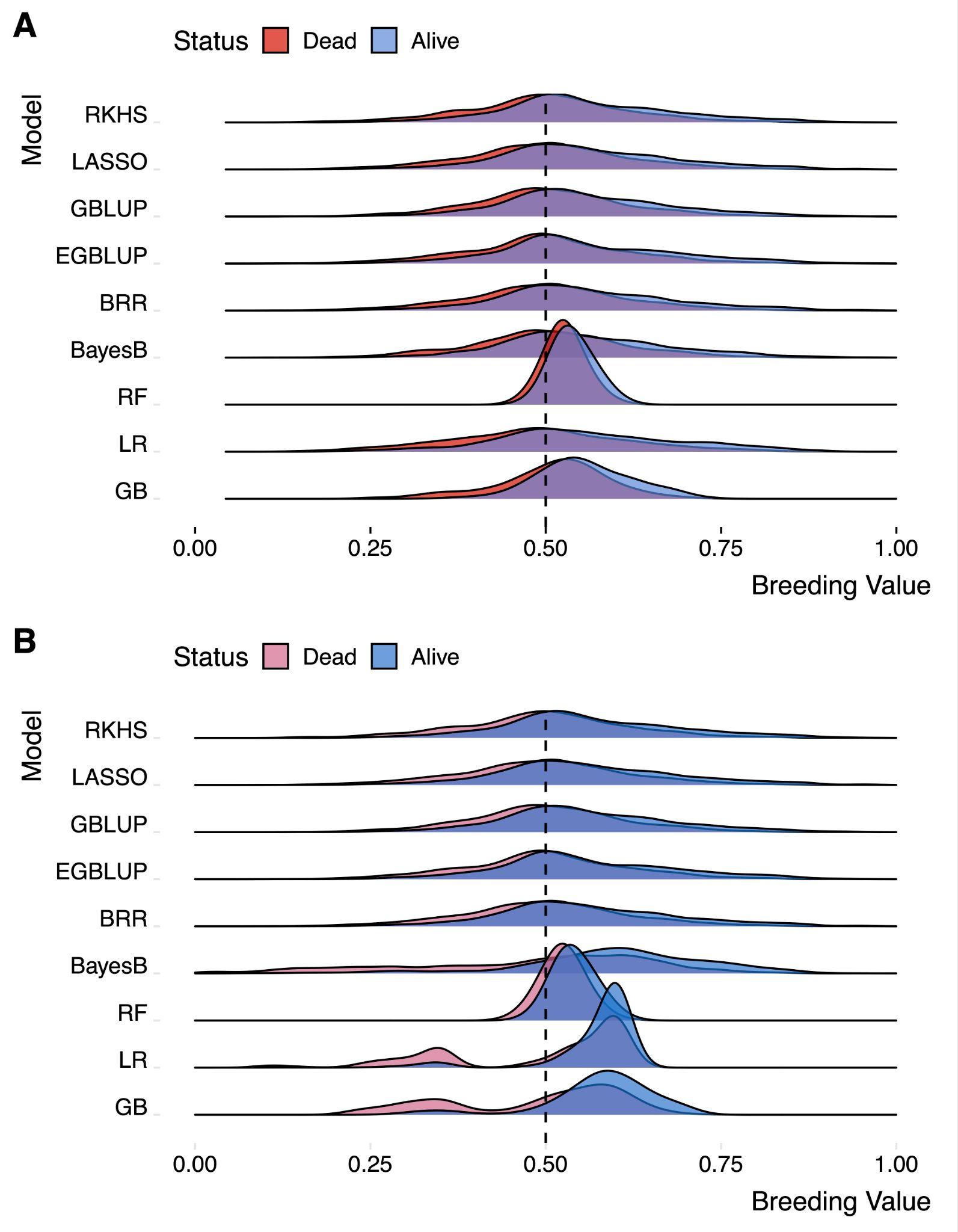

**Supplementary Figure 6:** Distribution of genome-estimated breeding values (GEBVs) **(A)** before and **(B)** after the addition of GSMs to the training dataset at MAF = 0.05. The black dashed line indicates a breeding value of 0.5.

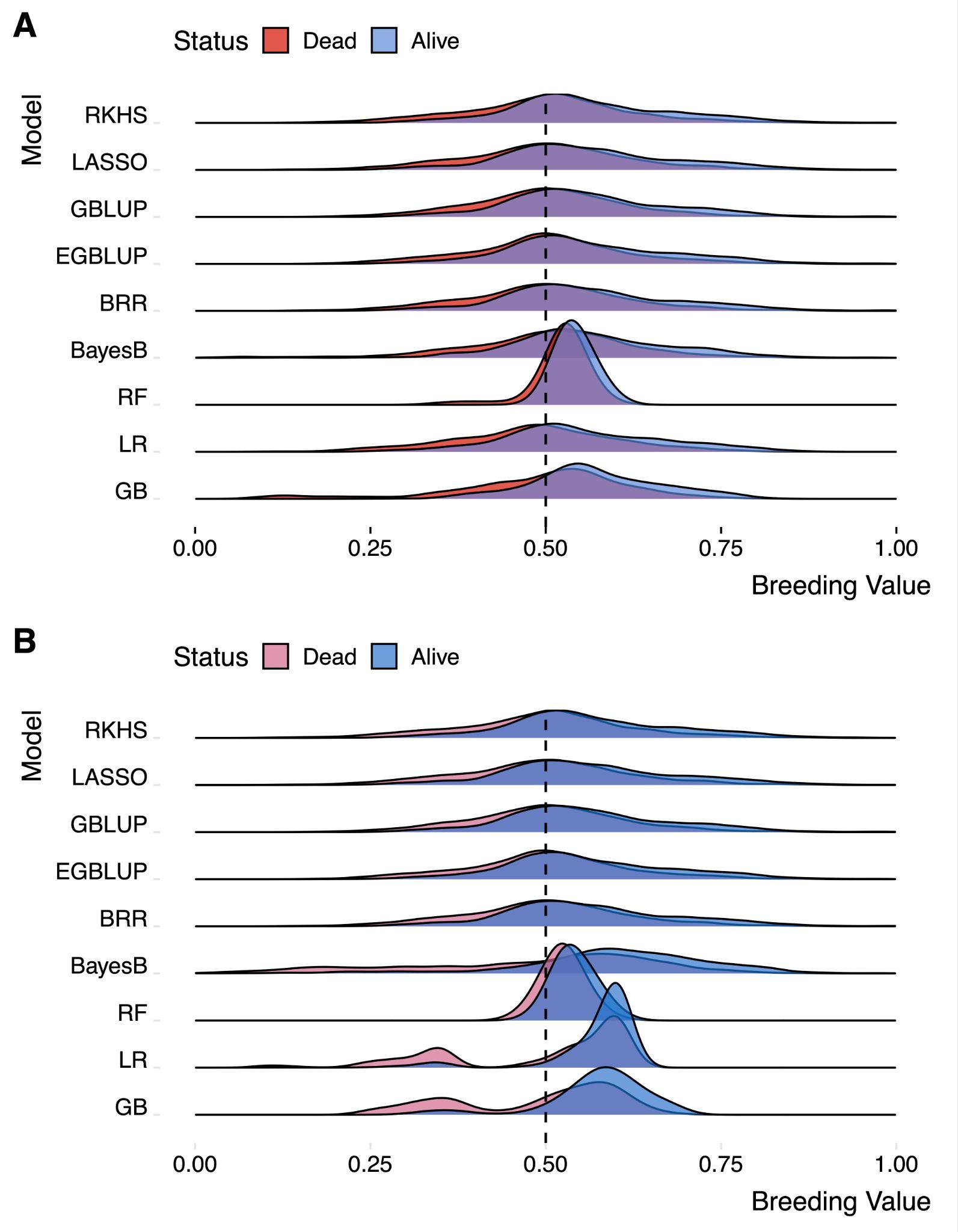

**Supplementary Figure 7:** Distribution of genome-estimated breeding values (GEBVs) **(A)** before and **(B)** after the addition of GSMs to the training dataset at MAF = 0.01. The black dashed line indicates a breeding value of 0.5.
